# K63-linked ubiquitin chains mark inactive Smoothened for packaging into ciliary extracellular vesicles

**DOI:** 10.64898/2026.07.01.735856

**Authors:** Mingli Zhu (祝明莉), Yash Raj, Gary A. Bradshaw, David U. Mick, Marian Kalocsay, Maxence V. Nachury

**Affiliations:** Department of Ophthalmology, University of California San Francisco; CA 94143, USA; Smith Cardiovascular Research Institute, University of California San Francisco; CA 94143, USA; Department of Systems Biology, Laboratory of Systems Pharmacology, Harvard Medical School; Boston, MA 02115, USA; Center for Molecular Signaling, Department of Medical Biochemistry and Molecular Biology, Saarland University School of Medicine; Homburg, Germany; Department of Experimental Radiation Oncology, The University of Texas MD Anderson Cancer Center; Houston, TX 77030, USA

## Abstract

Signaling receptors can exit cilia either by retrieval back into the cell or by secretion in extracellular vesicles (EVs), a process known as ectocytosis. The mechanisms that govern ectocytosis remain poorly understood. Here, we leverage quantitative proteomic profiling to identify the Hedgehog signaling receptor Smoothened (SMO) as a major cargo of cilia-derived EVs. Surprisingly, ligands that promote ciliary accumulation of SMO strongly suppressed its packaging into EVs, indicating that ectocytosis selectively packages specific conformational states of SMO. We further find that SMO packaged into EVs is extensively modified with K63-linked ubiquitin chains. Preventing SMO ubiquitination or selectively removing K63-linked ubiquitin chains from SMO markedly reduced its secretion into EVs and caused SMO accumulation within cilia. Together, these results identify K63-linked ubiquitin chains as a sorting signal for ciliary ectocytosis and indicate that ubiquitin-dependent packaging of inactive SMO contributes to the dynamic redistribution of SMO during Hedgehog signaling.

## INTRODUCTION

Cilia are specialized, surface-exposed organelles that play a pivotal role in organizing and transducing signaling pathways^1–3^. Central to Hedgehog (Hh) signaling is the regulated accumulation of the G protein-coupled receptor (GPCR) Smoothened (SMO) in cilia^4^. One mechanism controlling ciliary SMO abundance is BBSome-mediated retrieval, in which the intraflagellar transport (IFT) machinery and the BBSome complex return membrane proteins from cilia back into the cell. In unstimulated cells, SMO continuously enters and exits cilia, and single-molecule imaging studies indicate that this constitutive exit is mediated by the BBSome^5,6^.

Activation of the Hh pathway suppresses SMO retrieval, thereby promoting its accumulation in cilia^5–7^. The regulated attachment of K63-linked ubiquitin (K63Ub) chains, a modification otherwise associated with endolysosomal sorting^8^, marks GPCRs for recognition by the ubiquitin readers TOM1L2 and CFAP36 and subsequent removal from cilia by the BBSome and the IFT machinery^9–13^.

An alternate modality for activity-dependent exit of GPCRs from cilia is through their packaging into extracellular vesicles (EVs) that bud from the ciliary membrane, a process known as ciliary ectocytosis^14–16^. This evolutionarily conserved process not only serves to dispose of excess ciliary material when the capacity of retrieval is exceeded^17–20^ but may also facilitate intercellular communication in processes such as mating^21,22^, chemotaxis^23^, and immune evasion^24^. However, a major obstacle in testing the physiological importance of ciliary ectocytosis is that its underlying mechanisms remain largely uncharacterized. Thus, whether ciliary ectocytosis contributes to the dynamic regulation of SMO abundance in cilia remains unknown.

More fundamentally, we do not understand how the ectocytosis machinery selects cargoes for packaging into ciliary EVs. Because cargo selection in membrane trafficking is often dictated by post-translational modifications, ubiquitination represents an attractive candidate sorting signal. Although ubiquitination is a well-established sorting signal for BBSome-mediated retrieval from cilia, whether ubiquitin also directs cargo selection into ciliary EVs remains unknown.

In this study, we identify SMO as a major cargo of ciliary EVs in mammalian cells and demonstrate that K63-linked ubiquitin chains are required for its packaging into EVs. We further show that ectocytosis selectively packages a signaling-incompetent form of SMO. These findings reveal an unexpected mechanistic overlap between ectocytosis and BBSome-mediated retrieval and suggest that ectocytosis contributes to the dynamic redistribution of SMO during Hedgehog signaling.

## RESULTS

### Proteomic Profiling of Ciliary EVs Uncovers Candidate Cargoes and Regulators of Ectocytosis

To gain insight into the mechanisms of signal-dependent ciliary ectocytosis, we set out to inventory the composition of ciliary EVs via quantitative proteomics. We chose mouse IMCD3 cells as a model system due to their ability to accurately recapitulate ciliary trafficking and signaling mechanisms. Consistent with previous studies^17^, ARL13B, a marker of the ciliary membrane, was recovered in the small EV (sEV) fraction by differential ultracentrifugation, and genetic ablation of cilia in *Cep164^-/-^* cells removed ARL13B from the sEV fraction without affecting the secretion of the broad EV marker CD9 (**Fig. S1A**). A first approach to identify proteins secreted via cilia into EVs relied upon the Cilia-APEX2 system^25,26^ to selectively biotinylate proteins within cilia prior to EV production and isolation of biotinylated proteins from sEVs (**Fig. S1B**). This approach identified 129 proteins, including several subunits of intraflagellar transport complex B (IFT-B), the IFT motor kinesin-2 and the BBSome.

A second approach based on protein correlation profiling took advantage of the increase in ciliary ectocytosis when retrieval is compromised (such as in *Ift27^-/-^* cells) and its absence when cilia are ablated in *Cep164^-/-^* cells. Profiling the sEV proteomes from quadruplicate WT, *Cep164^-/-^*, and *Ift27^-/-^* samples by quantitative tandem mass tagging (TMT) mass spectrometry^27–29^ (**Fig. 1A**) revealed one cluster matching the expected signature of ciliary sEV (*Ift27^-/-^* ≫ WT > *Cep164^-/-^*) (**Figs. 1B and S1C**).

**Figure 1.**
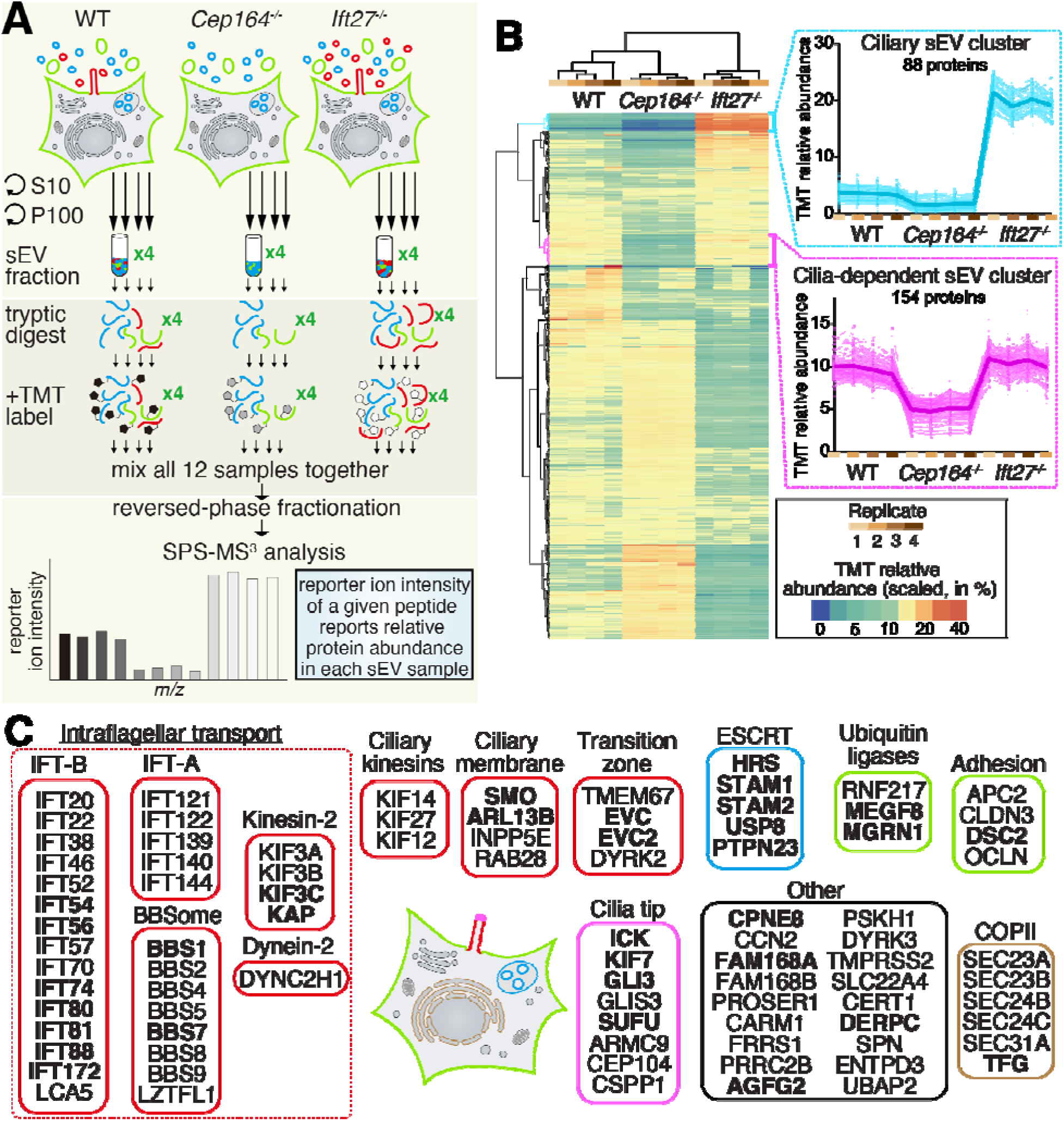
Quantitative proteomic profiling of ciliary EVs. **A**. Diagram of the proteomic profiling workflow. WT IMCD3 cells, ciliation-defective (*Cep164*^-/-^) cells, and retrieval-defective (*Ift27*^-/-^) cells were cultured to near-confluency and then serum-starved to induce ciliation and EV release for 20 h. EVs were purified from culture supernatants by differential ultracentrifugation ^39^. The small EV (sEV) fraction was lysed, cleaned on SP3 beads ^40^, and peptides were released by on-beads digest with trypsin. Each sEV sample was labeled with a unique tandem mass tag (TMT). All samples were then pooled and peptides were analyzed using a synchronous precursor selection MS^3^ method for mass spectrometric identification and quantitation. Each genotype was analyzed in biological quadruplicate. **B.** Two-way hierarchical cluster analysis of the sEV proteomes. The relative abundance of a given protein was calculated by dividing the TMT signal in one sample by the sum of the TMT signals in all samples. The color scheme of relative abundance is shown on the bottom right (in percent). All identified proteins are shown in the heatmap. The two sEV clusters related to cilia are shown in magnified view with each thin line representing an individual protein and the thick line representing the average of all proteins in the cluster. **C**. Heuristic representation of the ciliary sEV cluster. Proteins were categorized into subgroups based on their known localization and function. Hits in the Cilia-APEX2 sEV proteome are shown in bold.

The ciliary sEV cluster predominantly included known ciliary proteins, encompassing nearly all subunits of IFT-A, IFT-B, BBSome, and kinesin-2, the motor subunit of dynein-2, along with a few transition zone proteins and ciliary membrane proteins (**Fig. 1C**). Consistent with prior studies finding EV budding from the tip of cilia^14,15,30^, most proteins known to localize to the ciliary tip were found in the ciliary EV cluster. Over a third of the proteins in the ciliary sEV cluster were part of the Cilia-APEX2 sEV proteome including SMO, ARL13B, the SMO ubiquitin ligase complex MEGF8/MGRN1^31^, and the ESCRT-0 complex comprised of HRS and STAM or its close paralogue STAM2. Together, these proteomic approaches indicate that the ciliary sEV cluster represents a statistically validated proteome of EVs secreted via cilia in IMCD3 cells.

In addition to the ciliary sEV signature profile (*Ift27^-/-^*≫ WT > *Cep164^-/-^*), the sEV proteome profiling approach revealed one cluster of sEV proteins that were merely cilia-dependent (WT ∼ *Ift27^-/-^*> *Cep164^-/-^*) (**Figs. 1B and S1C**). In contrast to the ciliary sEV cluster that consisted of 55% of known ciliary proteins and 36% of proteins identified in the Cilia-APEX2 sEV proteome, fewer than 3% of proteins in the cilia-dependent sEV cluster were known ciliary proteins, and fewer than 6% were present in the Cilia-APEX2 sEV proteome. These results suggest that these proteins from the cilia-dependent sEV cluster are not secreted via cilia, even though their secretion may be cilium-dependent. Notably, the cilia-dependent sEV cluster included many subunits from the ESCRT-I and ESCRT-III/VPS4 complexes. The existence of cilia-dependent sEVs that may be secreted via non-ciliary routes underscores the importance of combining multiple cilia-less and retrieval-defective cells to define the proteome of ciliary sEVs.

### SMO Ectocytosis is Regulated by its Conformational State

To validate our mass spectrometry findings, we generated cell lines stably expressing moderate levels of SMO fused to either FLAG or NanoLuciferase (NLuc) (**Fig. S2A**) and verified that the signal- and BBSome-dependent ciliary dynamics of SMO^FLAG^ and SMO^NLuc^ (**Figs. S2B,** see also **Fig. 3F and H**) mirrored those of endogenous SMO^32^. We immunoblotted sEVs from WT, *Cep164^-/-^*, and *Ift27^-/-^* cells for SMO^FLAG^ or SMO^NLuc^, ARL13B, the generic EV marker CD9 and the ER marker GP96 (**Figs. 2A and S2C**). Measurement of the secretion ratio (the total amount detected in sEVs normalized to the total amount in cells, see methods) revealed that approximately 2% of SMO^FLAG^ or SMO^NLuc^ is secreted into sEVs. In agreement with our quantitative proteomics, the secretion of both SMO and ARL13B, but not that of CD9, was strictly cilium-dependent and the secretion ratio increased to nearly 5% when retrieval was compromised. These results indicate that SMO is efficiently selected for packaging into ciliary EVs.

**Figure 2.**
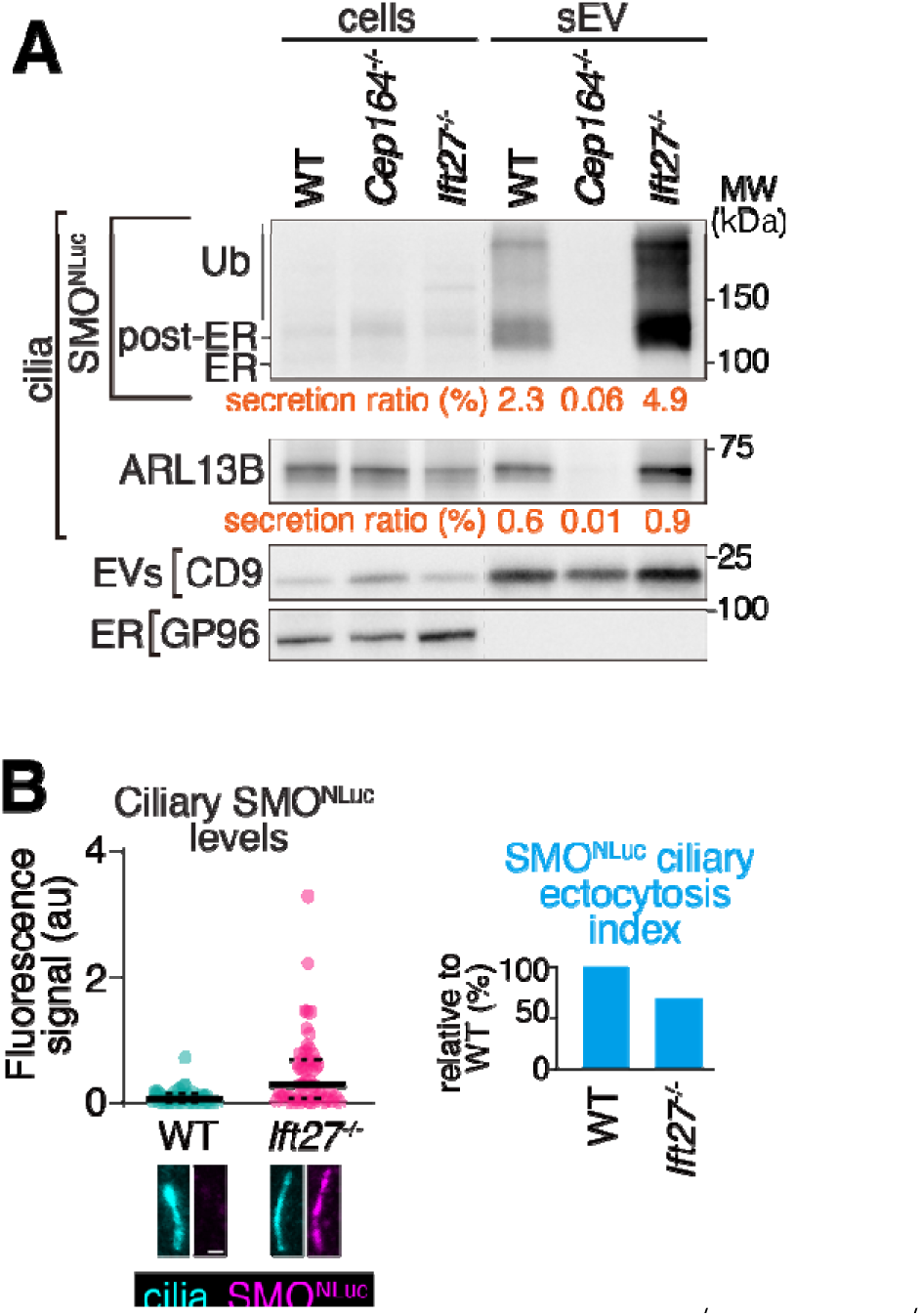
SMO is a major cargo of ciliary EVs. Near-confluent WT, *Cep164*^-/-^, and *Ift27*^-/-^ IMCD3 cells stably expressing SMO-FLAG-NLuc (SMO^NLuc^) were serum-starved and EV secretion allowed to proceed for 20 h. **A.** sEV were prepared from the culture supernatants and cell lysates and sEV fractions were immunoblotted for FLAG, ARL13B, CD9, and GP96. The secretion ratios of SMO^NLuc^ and ARL13B are shown below the immunoblots. 10 μg of cell lysates (corresponding to 0.1-0.19% of the total samples) were loaded for all immunoblots. 30% of the P100 fractions were loaded for SMO and ARL13B detection, and 10% for CD9 and GP96 detection. **B.** Cells were stained for FLAG and acetylated tubulin (cilia) and the ciliary fluorescence intensity in the FLAG channel was plotted in a dot plot. In this and every dots plot, the thick bar indicates the median and the dotted lines the first and third quartiles. *N* = 50 cilia were analyzed for each cell line; au, arbitrary units. Representative cilia images shown below the graph. Scale bar, 1 μm. The ciliary ectocytosis indices of SMO^NLuc^ were plotted in a bar graph.

**Figure 3.**
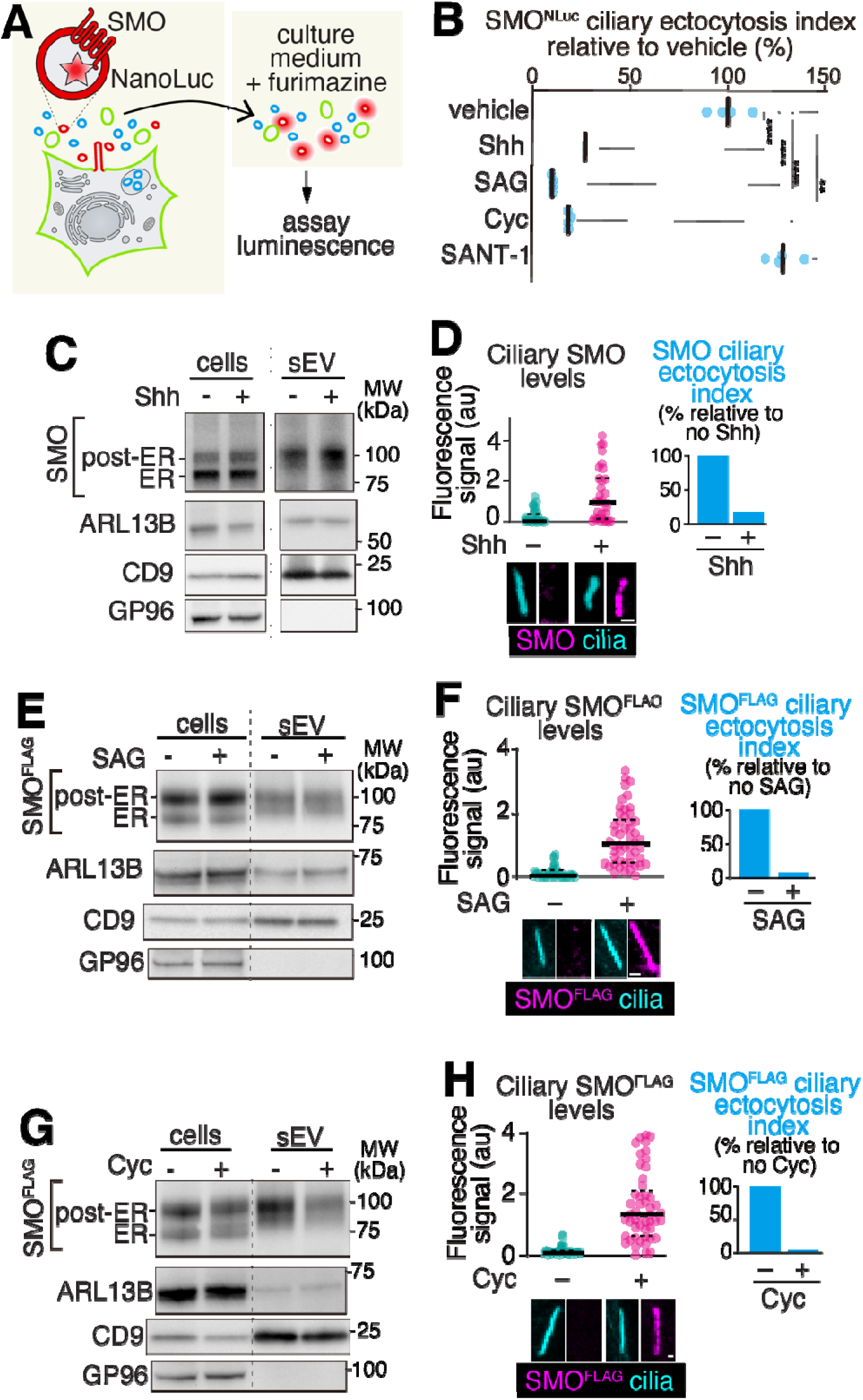
The inactive form of SMO is secreted into EVs via cilia. **A.** Diagram of the NanoLuciferase SMO secretion assay. IMCD3-[SMO-FLAG-NLuc] cells were cultured to near-confluency and then serum-starved to induce ciliation and release of ciliary EVs. Luciferase activity was measured in culture supernatants and cells with the Nano-Glo lucif rase assay. **B.** Regulation of SMO^NLuc^ secretion by signaling. Near-confluent IMCD3-[SMO-FLAG-NLuc] cells were serum-starved for 20 h. Fresh starvation medium supplemented with 20% Shh conditioned medium, 200 nM SAG, 5 μM cyclopamine, 200 nM SANT-1, or vehicle was added to cells and EV production was allowed to proceed for 4 h. The levels of SMO^NLuc^ in culture supernatants and in cells were measured using the NanoLuc assay. In parallel, identically treated cells grown on coverslips were fixed and stained to measure the ciliary SMO^NLuc^ levels. Each blue dot represents the SMO^NLuc^ ciliary ectocytosis index from one experiment. *N* = 4 independent experiments. Asterisks indicate unpaired *t*-test significance values. **** *p* < 0.0001; ** *p* < 0.005. **C-D.** Regulation of endogenous SMO secretion by Shh. Near-confluent NIH-3T3 cells were serum-starved and treated with or without 20% Shh-conditioned medium for 20 h. **C.** sEVs were prepared from the culture supernatants, and cell lysates and sEV fractions were immunoblotted for SMO, ARL13B, CD9, and GP96. See **Fig. S2F** for validation of the anti-SMO antibody. 20 μg of cell lysates (corresponding to 0.3% of the total sample) were loaded for all immunoblots. 20% of the P100 fractions were loaded for SMO and ARL13B detection, and 10% for CD9 and GP96 detection. **D.** Cells were stained for SMO and acetylated tubulin (cilia) and the ciliary fluorescence intensity in the SMO channel was plotted in a dot plot. *N* = 30-36 cilia analyzed per condition. Representative cilia images shown below the graph. Scale bar, 1 μm. The ciliary ectocytosis indices of SMO were plotted in a bar graph. **E-F.** Regulation of SMO^FLAG^ secretion by SAG. Near-confluent IMCD3-[SMO-FLAG] cells were serum-starved and treated with or without 200 nM SAG for 20 h. **E.** sEVs were prepared from the culture supernatants, and cell lysates and sEV fractions were immunoblotted for FLAG, ARL13B, CD9, and GP96. 0.2% of the total cell lysates were loaded for all immunoblots. 30% of the P100 fractions were loaded for SMO and ARL13B detection, and 10% for CD9 and GP96 detection. **F.** Cells were stained for FLAG and acetylated tubulin (cilia) and the ciliary fluorescence intensity in the FLAG channel was plotted in a dot plot. *N* = 50 cilia analyzed per condition. Representative cilia images shown below the graph. Scale bar, 1 μm. The ciliary ectocytosis indices of SMO^FLAG^ were plotted in a bar graph. **G-H.** Regulation of SMO^FLAG^ secretion by cyclopamine. Near-confluent IMCD3-[SMO-FLAG] cells were serum-starved and treated with or without 5 μM cyclopamine (Cyc) for 20 h. Immunoblots (**G)** were conducted as in **E** and cell staining, fluorescence measurements and ciliary ectocytosis indices **(H)** were performed and calculated as in **F**.

We next asked whether the increased SMO secretion in the *Ift27^-/-^* retrieval mutant is simply caused by the increased amounts of SMO in cilia when retrieval is blocked. If all the molecules of SMO inside cilia are equally likely to get packaged into EVs, the amounts of SMO secreted into EV will scale with the amount of SMO in cilia. To determine the likelihood that a given SMO molecule in cilia becomes packaged into EV, we normalized the amount of SMO detected in EVs to the fluorescent signal of SMO inside cilia (ciliary ectocytosis index). The ectocytosis index of SMO^FLAG^ and SMO^NLuc^ was similar in *Ift27^-/-^* and WT cells (**Figs. 2B and S2D**), indicating that forced ciliary accumulation of SMO does not intrinsically affect its ectocytosis efficiency.

The relatively high flux of SMO ectocytosis suggested that this process may participate in the signal-dependent trafficking of SMO. To streamline the measurements of SMO secretion under different conditions, we directly detected SMO^NLuc^ in culture supernatants and cells using luminescence (**Fig. 3A**). The luciferase assay faithfully reported on ciliary ectocytosis, as SMO^NLuc^ secretion was reduced more than 5-fold when cilia were removed in *Cep164^-/-^*cells (**Fig. S2E**). To determine whether SMO secretion is regulated by its activity, we treated cells with the natural ligand Sonic Hedgehog (Shh), the SMO agonist SAG, or the SMO antagonists cyclopamine and SANT-1. The ectocytosis index of SMO^NLuc^ was reduced by 75 to 90% when cells were treated with cyclopamine, SAG, or Shh, and modestly increased with SANT-1 treatment (**Fig. 3C**).

Notably, cyclopamine decouples ciliary accumulation from activation, as this SMO ligand blocks Hedgehog pathway activity while promoting SMO accumulation inside cilia ^33,34^. Thus, all the ligands that promoted ciliary enrichment of SMO, regardless of their ability to activate SMO, decreased the likelihood of SMO^NLuc^ becoming packaged into EVs. We validated these results with conventional EV preparations and endogenous SMO in 3T3 cells, a well-accepted cell line for studying the Hh response^35^. Here, activation of the Hedgehog pathway with Shh reduced the ciliary ectocytosis index of endogenous SMO by 82%, confirming that SMO activation by natural means drastically suppresses its packaging into EVs at cilia (**Fig. 3C-D**).

We tested the influence of SAG and cyclopamine on SMO^FLAG^ secretion with conventional EV preparations in the IMCD3-[SMO-FLAG] cell line and found that both ligands, which promote ciliary accumulation despite having opposite effects on its downstream signaling, decreased the ciliary ectocytosis index of SMO by more than 92% (**Fig. 3E-I**). We note that the secretion of ARL13B remained unaffected by treatment with either cyclopamine (**Fig. 3E**), SAG (**Fig. 3G**) or Shh (**Fig. 3I**). Similarly, other proteins from the ciliary sEV cluster, including EVC, HRS, and STAM2, showed no significant changes in their secretion ratio in response to SAG treatment (**Fig. S2G**).

The profound decrease in the efficiency of ciliary ectocytosis when SMO is bound to ligands that promote its ciliary retention contrasts with the unchanged efficiency of ciliary ectocytosis of SMO in *Ift27^-/-^* cells (**Figs. 2B and S2D**). These results indicate that acquisition of a conformation competent for ciliary enrichment strongly reduces SMO’s likelihood of being packaged into sEVs at cilia. We conclude that the ciliary ectocytosis machinery preferentially selects SMO molecules that have yet to adopt a conformation competent for ciliary retention.

### K63-linked Ubiquitin Chains are Required for SMO Secretion into sEVs

We noted that a significant proportion of SMO in the sEV fraction appeared as a high molecular weight smear, indicative of poly-ubiquitination (**Figs. 2A and S2C**). Indeed, immunoprecipitation of SMO from the sEV fraction revealed abundant K63-linked ubiquitin chains associated with SMO (**Fig. 4A**). Furthermore, treatment of sEV lysates with either USP2 (a broad spectrum deubiquitinase) or AMSH (a deubiquitinase specific for K63-linkages) reduced the SMO smear to a similar extent (**Fig. 4B**), indicating that K63 linkages predominate in the ubiquitin chains attached to SMO in sEVs.

**Figure 4.**
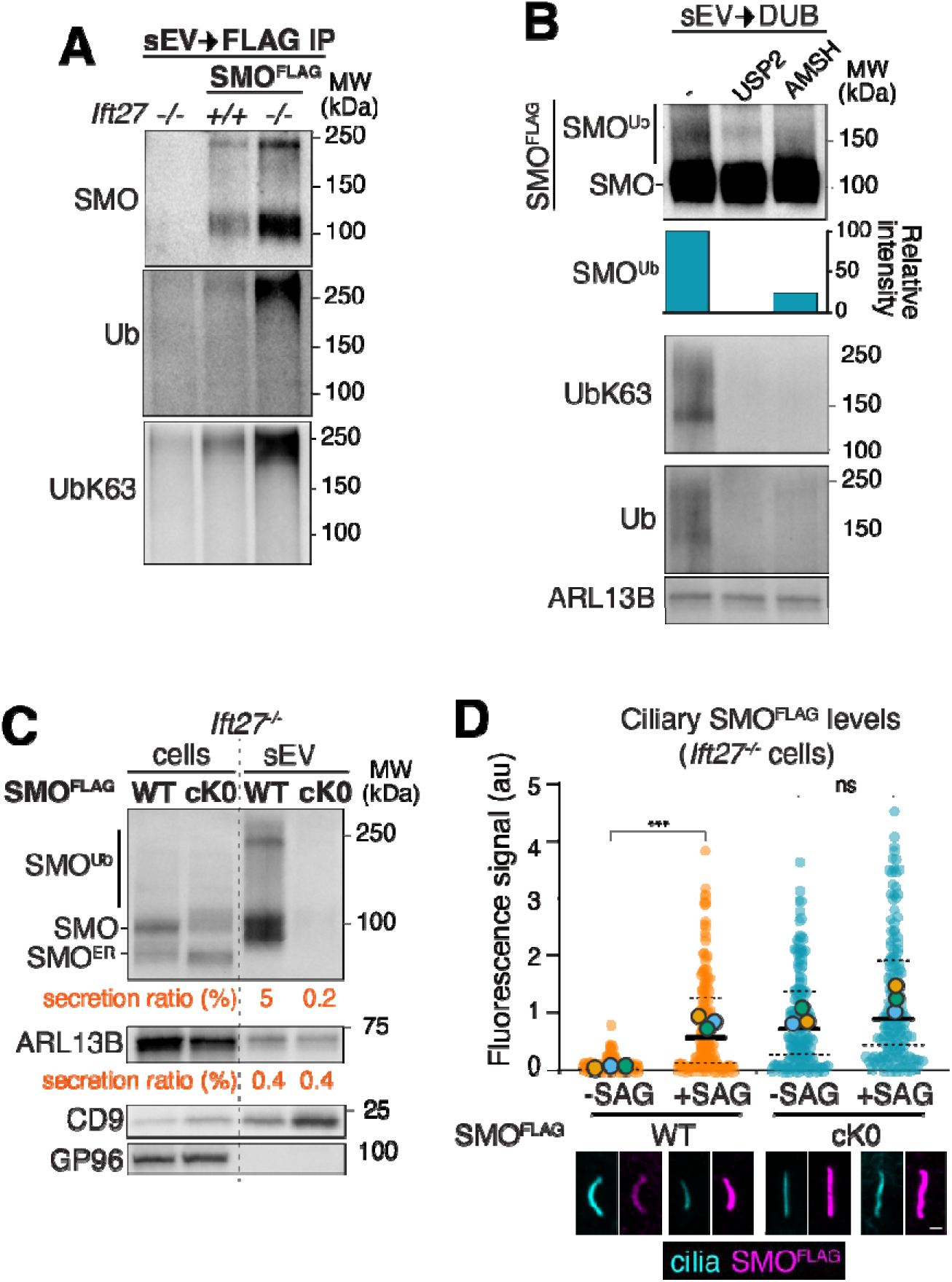
SMO ubiquitination is required for its secretion into sEV. **A.** SMO ubiquitination in EVs. sEVs prepared from near-confluent, serum-starved *Ift27*^-/-^ IMCD3 cells and WT and *Ift27*^-/-^ IMCD3-[SMO-FLAG] cells were lysed, SMO^FLAG^ was captured on anti-FLAG resin and eluates were immunoblotted for FLAG, Ub, and UbK63 chains. **B.** Deubiquitinase treatment of sEV. sEVs prepared from *Ift27*^-/-^ IMCD3-[SMO-FLAG] cells were lysed and treated with 2 μM USP2 or 2 μM AMSH. Samples were immunoblotted for FLAG, UbK63, Ub, and ARL13B. The relative intensity of the high-molecular weight smear of SMO^FLAG^ was quantitated and shown. **C.** Effect of cytoplasmic lysine removal (cK0) on SMO secretion. sEVs were prepared from *Ift27*^-/-^ IMCD3 cells stably expressing either SMO^FLAG^ or SMO^FLAG^ cK0. Cell lysates and sEV fractions were immunoblotted for FLAG, ARL13B, CD9, and GP96. 20 μg of cell lysates (corresponding to 0.42-0.55% of the total sample) were loaded for all immunoblots. 25% of the P100 fractions were loaded for SMO and ARL13B detection, and 10% for CD9 and GP96 detection. The secretion ratios of SMO^FLAG^ and ARL13B and the ciliary ectocytosis indices of SMO^FLAG^ were quantified and are shown. **D.** Effect of cytoplasmic lysine removal (cK0) on SMO levels in cilia. *Ift27*^-/-^ IMCD3 cells stably expressing SMO^FLAG^ or SMO^FLAG^ cK0 mutant were treated with or without 200nM SAG and stained for FLAG and acetylated tubulin (cilia) and the ciliary fluorescence intensity in the FLAG channel was plotted in a dot plot. *N* = 146-163 cilia from 3 independent experiments; Asterisks indicate unpaired *t*-test significance values. ns, non-significant (*p* = 0.10). ***, *p* < 0.001. au, arbitrary units. Representative cilia shown below the graph. Scale bar, 1 μm.

K63-linked ubiquitin chains instruct sorting decisions in membrane trafficking, primarily targeting membrane proteins for lysosomal degradation, but also marking GPCRs for retrieval from cilia^9,10^. To determine whether ubiquitination marks SMO for packaging into ciliary EVs, we eliminated SMO ubiquitination by mutating all cytoplasmic lysine residues of SMO to arginine (cK0). SMO^cK0^’s secretion ratio was reduced more than 20-fold compared to wild-type SMO (**Fig. 4C**), without affecting ARL13B or CD9 secretion. Correspondingly, blocking SMO ubiquitination increased ciliary SMO levels by nearly 10-fold in the *Ift27^-/-^*retrieval mutant, with ciliary SMO^cK0^ levels reaching the near-maximal levels seen after treatment with the SMO agonist SAG (**Fig. 4D**).

To specifically test the role of K63Ub linkages in sorting SMO into EVs, we fused the catalytic domain of AMSH to SMO (**Fig. 5A**). AMSH reduced SMO secretion by 5- to 10-fold, while the catalytically inactive AMSH left SMO secretion unchanged (**Fig. 5B**). Again, the blockage of SMO ectocytosis resulted in a dramatic increase in ciliary SMO levels (**Fig. 5C**) with ciliary SMO-AMSH reaching levels normally seen in SAG-treated cells (**Fig. S3**). Blocking retrieval did not significantly increase the ciliary levels of SMO-AMSH (**Figs. 5C and S3**), indicating that removal of UbK63 chains from SMO blocks both ectocytosis and retrieval. These results collectively demonstrate that K63Ub chains mark SMO for packaging into ciliary EVs.

**Figure 5.**
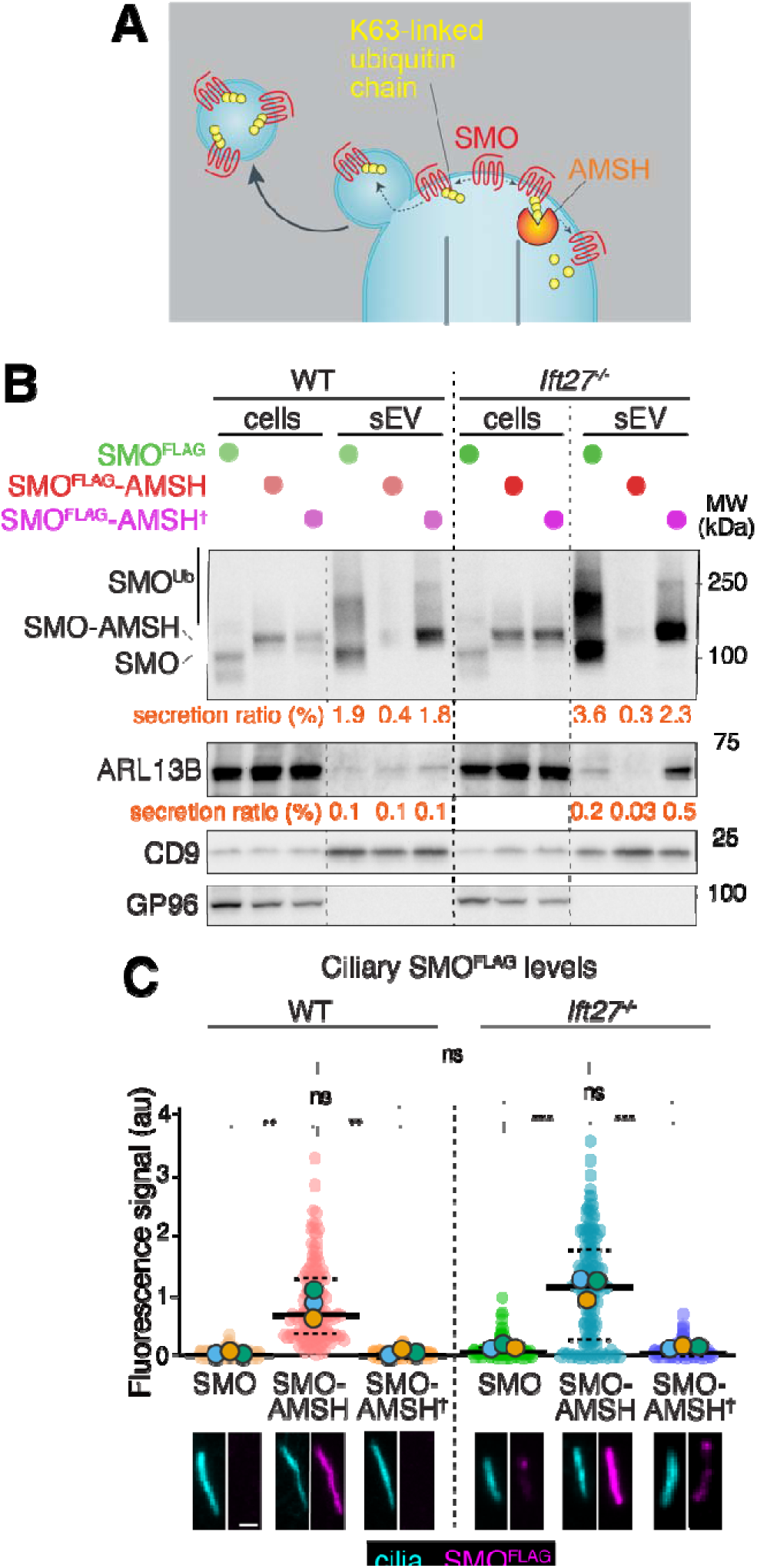
UbK63 chains are required for SMO secretion into sEV. **A.** Diagram of the targeted deubiquitination strategy. **(B-C)** Effect of forced deubiquitination of SMO ectocytosis. **B.** sEVs were prepared from WT and *Ift27*^-/-^ IMCD3 cells stably expressing SMO^FLAG^, SMO^FLAG^-AMSH, SMO^FLAG^-AMSH (catalytically inactive E38A mutant). Cell lysates and sEV fractions were immunoblotted for FLAG, ARL13B, CD9, and GP96. For SMO and ARL13B detection, 20 μg of cell lysates (corresponding to 0.36-0.76% of the total sample) and 20% of the P100 fractions were immunoblotted. For CD9 and GP96 detection, 10 μg of cell lysates (corresponding to 0.18-0.38% of the total sample) and 5% of the P100 fractions were immunoblotted. The secretion ratios of SMO^FLAG^ and ARL13B and the ciliary ectocytosis indices of SMO^FLAG^ were quantified and are shown. **C.** Cells were stained for FLAG and acetylated tubulin (cilia) and the ciliary fluorescence intensity in the FLAG channel was plotted in a dot plot. *N* = 150 cilia from 3 independent experiments; Asterisks indicate unpaired *t*-test significance values. ns, non-significant (*p* = 0.201). **, *p* < 0.01, ***, *p* < 0.001. au, arbitrary units. Representative cilia images shown below the graph. Scale bar, 1 μm.

## DISCUSSION

### K63-linked ubiquitin chains act as a sorting signal for ciliary ectocytosis

Our findings identify K63-linked ubiquitin chains as a sorting signal for ciliary ectocytosis. While ubiquitination is known to participate in the biogenesis of ARRDC1-mediated microvesicles^36–38^ and other membrane trafficking events, its role in the selection of cargoes for ciliary EVs had remained unknown. We find that SMO molecules packaged into ciliary EVs are heavily modified with K63-linked ubiquitin chains and that preventing SMO ubiquitination or selectively removing UbK63 chains from SMO markedly suppresses its secretion into EVs. Together, these results establish ubiquitination as a molecular signal that earmarks cargoes for ciliary ectocytosis.

The identification of UbK63 chains as a sorting signal for ectocytosis is particularly notable because the same modification directs GPCR retrieval from cilia^9,11^. Thus, cargo selection for the two major exit pathways from cilia relies on a common molecular mark. This finding reveals an unexpected mechanistic overlap between retrieval and ectocytosis. While our finding that UbK63 chains act as a sorting signal for ectocytosis help explain why ciliary GPCRs fused to Ub disappeared from cilia of retrieval-defective cells^11^, the discovery that the same post-translational modification promotes sorting into ciliary EVs and retrieval presents a conundrum.

Several models may explain how a common ubiquitin signal feeds two distinct exit pathways. Retrieval and ectocytosis may compete for the same pool of ubiquitinated cargoes using distinct ubiquitin readers. In this scenario, retrieval would normally remove most ubiquitinated proteins from cilia, whereas ectocytosis would become favored when the local concentration of ubiquitinated cargoes exceeds the capacity of retrieval.

Alternatively, additional features of the ubiquitin modification itself, such as chain architecture, chain length, or associated binding partners, may bias cargoes toward one pathway or the other. Finally, other modifications besides ubiquitination may route cargoes to one route. Distinguishing among these possibilities will require identification of the molecular machinery that recognizes ubiquitinated cargoes during ciliary ectocytosis.

### Regulated ubiquitination may underlie selective cargo recognition during ectocytosis

Previous studies established that ciliary ectocytosis can selectively remove signaling receptors from cilia^17^. Our findings now suggest a molecular mechanism that may underlie this selectivity. We find that ligands promoting ciliary retention of SMO, regardless of whether they activate or inhibit downstream Hedgehog signaling, strongly suppress SMO packaging into EVs. Thus, ectocytosis does not simply track the abundance of SMO within cilia. Instead, it preferentially selects a subset of SMO molecules that have not acquired a conformation competent for ciliary retention.

Our studies identify SMO as a major cargo of ciliary EVs in IMCD3 and NIH-3T3 cells, with up to 5% of the total cellular SMO pool secreted into ciliary EVs over a 16-hour period. Combined with the preferential packaging of inactive SMO, this substantial flux suggests that ectocytosis contributes meaningfully to the selective removal of non-retained SMO molecules from cilia.

SMO is proposed to exist in a dynamic equilibrium between conformations that differ in their ability to reside within cilia^33,34^. We find that ligands promoting ciliary retention, whether in an active form (Shh, SAG) or inactive form (cyclopamine), lead to a pronounced suppression of SMO ectocytosis from cilia. The preferential packaging of cilia-incompetent SMO therefore suggests that ectocytosis contributes to the dynamic redistribution of SMO during Hedgehog signaling. More broadly, these findings raise the possibility that selective ubiquitination provides a biochemical mechanism for distinguishing among different conformational states of the same receptor and coupling this distinction to cargo sorting into EVs.

One proposed mechanism for regulated ubiquitination of SMO inside cilia involves the ubiquitin ligase WWP1^10^. This ligase is associated with the Hedgehog receptor Patched1, and Hedgehog-dependent removal of Patched1 from cilia decreases SMO ubiquitination. However, this model does not readily explain the effects of cyclopamine or SAG, both of which promote ciliary accumulation of SMO through direct actions on SMO rather than through the canonical inhibition and removal of Patched1 from cilia. These observations argue for the existence of ubiquitination machinery capable of directly recognizing SMO molecules that are not competent for ciliary retention.

In this context, the presence of the transmembrane ubiquitin ligase complex MEGF8/MGRN1 within the ciliary EV proteome is notable. MEGF8/MGRN1 promotes SMO ubiquitination and removal from the cell surface^31^ and therefore represents a plausible candidate for marking inactive SMO molecules for removal from cilia.

Determining how the ubiquitination machinery distinguishes among different conformational states of SMO will be essential for understanding how ciliary membrane composition is actively maintained.

### Concluding remarks

Together, our findings identify SMO as a major cargo of ciliary EVs and establish K63-linked ubiquitin chains as a sorting signal for its packaging into EVs. The substantial flux of SMO ectocytosis and the preferential packaging of inactive SMO suggest that ectocytosis contributes to the dynamic regulation of ciliary SMO abundance during Hedgehog signaling. More broadly, the shared requirement for UbK63 chains in retrieval and ectocytosis reveals an unexpected mechanistic connection between the two major exit pathways from cilia and raises the question of how ubiquitinated cargoes are routed between them. Resolving this question will provide important insights into how ciliary ectocytosis and the retrieval machinery select signaling receptors for removal from cilia.

## ACKNOWLEDGMENTS

We thank Maike De La Roche for the gifts of antibodies; Rohan Baker, David Komander, and Greg Pazour for the gifts of plasmids; Yien-Ming Kuo for help with microscopy; Lei Wang for use of the plate reader; Ryan Leib and the Stanford University Mass Spectrometry for data acquisition; Jaclyn Goldstein for help with the Cilia-APEX EV experiment; and all members of the Nachury lab for stimulating discussions. This work was funded by NIH (GM089933 to MVN) and ADA (1-20-VSN-03 to MVN). This work was made possible, in part, by EY002162 - Core Grant for Vision Research and by the Research to Prevent Blindness Unrestricted Grant (MVN).

The mass spectrometry proteomics data have been deposited to the ProteomeXchange Consortium via the PRIDE partner repository with the dataset identifier PXD057769.

## Materials and Methods

All reagents and resources used in this study are listed in Table S1.

### Plasmids

For bacterial expression, mouse SMO (aa 560-793) was amplified and cloned into pGPS1 (GST-HRV3C-Stag-) and pET-28(+) (6xHis-Thrombin-T7tag) by conventional cloning.

Coding sequences for FLAG-tagged mouse SMO (Addgene, plasmid no. 164532) and FLAG-tagged mouse SMO (cK0) (Addgene, plasmid no. 164545) were amplified by PCR and cloned in the vector pCrysB-FRT-DEST. NanoLuciferase (synthesized by GenScript), AMSH catalytic domain and the catalytically inactive mutant ^9^ were PCR amplified and cloned in the vector pCrysB-FRT-SMO-FLAG.

To knock out *Cep164* in IMCD3 Flp-In cells, a gRNA targeting exon 8 of mouse *Cep164* was cloned in pX459V2.0-eSpCas9(1.1) as described previously^25^.

### Cell culture

A parental IMCD3-FlpIn cell line (gift from Peter K. Jackson, Stanford University, Stanford, CA) was modified to generate all stable cell lines used in the study. IMCD3-FlpIn cells were cultured in DMEM/F12 (11330-057; Gibco) supplemented with 10% FBS (100-106; Gemini Bio-products), 100 U/mL penicillin-streptomycin (400-109; Gemini Bio-products), and 2 mM L-glutamine (400-106; Gemini Bio-products). Ciliation was induced by serum starvation in media with no FBS for 16 to 24 h.

The 3T3-FlpIn cell line (Thermo R76107) was cultured in DMEM (ThermoFisher, 11995-065) supplemented with 10% FBS, 100 U/mL penicillin-streptomycin, 2 mM L-glutamine and 1x non-essential amino acid (Cytiva SH30238.01). Ciliation was induced by serum starvation in media with no FBS for 16 to 24 h.

### Transfection and generation of stable cell lines

For the generation of stable cell lines that express SMO fusion proteins, a plasmid encoding the Flp recombinase (pOG44) was co-transfected with the FRT-based plasmids using XtremeGene9 (Roche) via reverse transfection method into IMCD3 Flp-In cells as described ^41^. Stable transformants were selected by blasticidin resistance (4 μg/mL).

For CRISPR-based genome editing of *Cep164* in IMCD3 Flp-In cells, Cas9 and guide RNA were transiently expressed from a pX459 derivative and transfectants selected with puromycin. Clones were isolated by limited dilution and screened for absence of cilia by immunofluorescence.

### Antibodies and drugs

The following mouse monoclonal antibodies were used for immunofluorescence: anti-acetylated tubulin (clone 6-11B-1; Sigma-Aldrich; 1:500), anti-FLAG (clone M2, F1804; Sigma-Aldrich; 1:1000).

The following rabbit polyclonal antibodies were used for immunofluorescence: anti-SMO (this study, 1:500).

The following monoclonal antibodies were used for immunoblotting: anti-FLAG (mouse; clone M2, F1804; Sigma-Aldrich; 1:1000), anti-ARL13B (mouse; ab136648; Abcam; 1:500), anti-CD9 (rat; sc-18869; Santa Cruz; 1:1000), anti-GP96 (rat; ADI-SPA-850-D; Enzo; 1:1000), anti-SMO (gift from Maike De La Roche; 1:500).

The following reagents were used at the indicated concentrations: 200 nM SAG, 5 μM cyclopamine, 200 nM SANT-1. Cyclopamine was dissolved in ethanol, SAG and SANT-1 were dissolved in DMSO.

### Microscopy

#### Fixed imaging

For fixed imaging, 50,000 cells were seeded on acid-washed 12 mm diameter #1.5 coverslips (12-545-81; Thermo Fisher Scientific), grown for 24 h, and then serum starved for 20 to 24 h before experimental treatment. After treatment, cells were fixed with 4% paraformaldehyde (50-980-487, Thermo Fisher Scientific) in PBS for 15 min at 37 °C. Cells were then permeabilized in IF buffer [PBS supplemented with 0.1% Triton X-100, 5% normal donkey serum (017-000-121; Jackson ImmunoResearch Laboratories), and 3% bovine serum albumin (BP1605-100; Thermo Fisher Scientific)]. Cells were then incubated at room temperature for 1 h with primary antibodies diluted in IF buffer, washed three times in IF buffer, and then incubated with secondary antibodies (Jackson ImmunoResearch Laboratories) diluted in IF buffer for 30 min. Cells were then washed three times with IF buffer and DNA was stained with Hoechst 33258 (H1398; Molecular Probes). Cells were washed twice more with PBS, and coverslips were mounted on slides using fluoromount-G (17984-25; Electron Microscopy Sciences).

Cells were imaged on confocal LSM 700 (Zeiss) or LSM 900 (Zeiss) microscopes equipped with 63x/1.4 NA oil objective, or on a Nikon AX NSPARC microscope with a 60x/1.4 NA oil objective. Z stacks were acquired at 0.5 μm interval.

### Image analysis

Files were imported from LSM700/900 or Nikon AX NSPARC workstations into ImageJ/ Fiji (National Institutes of Health) for all analyses.

### Measurement of ciliary signals

For the quantification of ciliary signals (for all the proteins, in all the figures) in fixed cells, maximum intensity projections were used. The ciliary intensities were measured using the following equation:

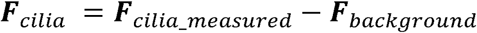

Where ***F****_cilia_measured_* is the total ciliary fluorescence detected in a mask that contains the entire cilium and ***F****_background_* is the fluorescence measured in an adjacent area ***F****_cilia_measured_* was measured in ImageJ by a mask consisting of a 3-pixel-wide line along the long axis of the cilium and an identical mask was used to measure ***F****_background_* in an adjacent area 2 to 3 pixels shifted to the side. For all measurements, the integrated density was used.

The ciliary intensities ***F****_cilia_* were plotted as dot plots using the GraphPad Prism. Each dot represents an individual data point, and all the data points are shown in the graphs. Median and interquartile range are marked by solid and dotted lines, respectively.

### EV methods

#### Small extracellular vesicle preparation

Small EV (sEV) were prepared as described^39^ with minor modifications. IMCD3 cells were cultured in DMEM/F12 medium supplemented with 10% FBS to near confluency in a 15 cm dish. The medium was then replaced with DMEM/F12 medium with no FBS added to induce ciliation and release of ciliary EVs. Medium was harvested 20-24 h later and extracellular vesicles were purified as diagramed in **Fig. 1A**. First, the culture supernatant was transferred to a 50 mL conical tube. The dishes were gently rinsed with 10 mL PBS, and the rinse was transferred to the conical tube. Second, the conical tube was centrifuged at 300 x *g* for 10 min at 4 □ to remove cell bodies. The supernatant (S3) was then transferred to a new 50 mL conical tube and centrifuged at 2000 x *g* for 20 min at 4 to remove cell debris (P2). The resulting supernatant (S2) was then centrifuged at 10,000 x *g* for 40 min at 4□ in a Beckman SW-32 swinging bucket rotor to pellet large EVs (P10). The resulting supernatant (S10) was then centrifuged at 100,000 x *g* for 90 min at 4□ in the SW-32 rotor. The pellet was washed in PBS and centrifuged at 100,000 x *g* for 90 min at 4□ in a Beckman SW-55 swinging bucket rotor. The final P100 pellet is the sEV fraction.

### Measurement of secretion ratio and ciliary ectocytosis index by by conventional EV purification and western blot

Purified sEVs (P100) and cell lysates were analyzed by immunoblotting. To calculate the secretion ratio of proteins, protein band intensities were measured in immunoblots of cell lysates and EV fractions, background intensities of an equivalent area in an empty lane were subtracted and the background-corrected intensities were scaled to the total fractions. The secretion ratio was calculated as follows:

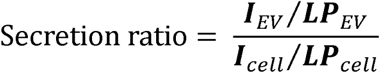

Where ***I*** is the background-corrected intensity of the protein band measured in the cell lysate or in the EV fraction; and ***LP*** is the proportion of each fraction loaded on the gel, i.e. the volume loaded on the gel divided by the total volume of the fraction.

To calculate the ciliary ectocytosis index of SMO, a coverslip was placed in the 15 cm plate used to produce the supernatant used in the EV preparation. The background-corrected band intensity was measured in the EV fraction immunoblotted for SMO and the average background-corrected ciliary level of SMO was measured in cells on the coverslip. The ciliary ectocytosis index was calculated as follows:

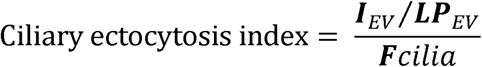

Where ***I*** is the background-corrected intensity of the protein band measured in the EV fraction; ***LP*** is the proportion of EVs loaded on the gel; and is the average background-corrected fluorescence intensity of SMO in cilia.

### Measurement of SMO^NLuc^ secretion by NanoLuc assays

IMCD3 cells stably expressing SMO^NLuc^ were cultured in DMEM/F12 supplemented with 10% FBS to near confluency in 96-or 24-well plates. The cells were then switched to DMEM/F12 with no FBS to induce ciliation and release of ciliary EVs. After 24 h, a NanoLuciferase assay was performed with a Nano-Glo® Luciferase Assay System (Promega N1110) according to the manufacturer’s instructions. As diagramed in **Fig. 3A**, starvation medium and cells were added to the reconstituted Nano-Glo® Luciferase Assay Reagent, and total luciferase activity was measured with a Multimode Plate Reader (BioTek Cytation 5).

### Measurement of secretion ratio and ciliary ectocytosis index by NanoLuc assays

To calculate the secretion ratio of SMO^NLuc^, the bioluminescence intensity of SMO^NLuc^ was measured in cell lysates and in culture supernatants by NanoLuc assays. The SMO secretion ratio was defined as the total bioluminescence intensity detected in starvation medium divided by the total bioluminescence intensity detected in cell lysate.

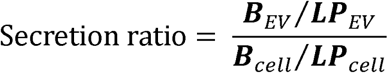

Where ***B*** is the bioluminescence intensity of SMO^NLuc^ measured in cell lysates and in culture supernatants by NanoLuc assays; and ***LP*** is the proportion of each fraction analyzed by NanoLuc assay.

We note that the SMO^NLuc^ secretion ratio measured by luciferase assay of culture supernatants (**Fig. S2E**) was higher than that the secretion ratio measured by immunoblotting of sEV fractions (**Fig. 2A**), likely reflecting the incomplete recovery of SMO-containing sEVs by purifying sEVs using differential ultracentrifugation.

To calculate the ciliary ectocytosis index of SMO^NLuc^, cells on coverslip were treated identically to the cells used to generate the culture supernatant analyzed by NanoLuc assays. The bioluminescence intensity of SMO^NLuc^ was measured in culture supernatants by NanoLuc assays and the average background-corrected ciliary level of SMO was measured in cells on the coverslip. The ciliary ectocytosis index was calculated as follows:

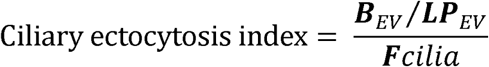

Where ***B*** is the bioluminescence intensity of SMO^NLuc^ measured in culture supernatants by NanoLuc assays; ***LP*** is the proportion of each fraction analyzed by NanoLuc assay; and is the average background-corrected fluorescence intensity of SMO in cilia.

### Production of Shh

Shh-containing conditioned medium was produced by HEK239T-EcR-Shh cells, which carry a stably integrated construct for expression of murine Shh under the control of an ecdysone-inducible promoter ^42^. Cells were grown to 80% confluence, the medium was changed to DMEM or DMEM/F12 with no FBS but supplemented with 5 μM ponasterone A, and conditioned medium was collected for 48 h and then filtered through a 0.22-μm filter (EMD Millipore). The titer of Shh was determined by testing dilutions of the conditioned media with NIH-3T3 Light-II reporter cells, which harbor stably integrated versions of GLI-driven firefly luciferase and constitutively expressed Renilla luciferase^35^. After stimulation, a dual-luciferase measurement was performed with Dual-Luciferase® Reporter Assay System (Promega E1960) according to the manufacturer’s instructions. Cells were lysed in Passive lysis buffer, and firefly luciferase activity and Renilla luciferase activity were measured with a Multimode Plate Reader (BioTek Cytation 5). As the minimum amount of conditioned medium needed for a saturating response was 10%, a concentration of 20% was used to maximally activate the pathway in **Figs. 3B-D and S2F**.

### Immunoprecipitation

*Ift27^-/-^* cells, WT cells and *Ift27^-/-^* cells stably expressing SMO^FLAG^ were cultured to near confluency and then serum-starved to induce ciliation and release of ciliary EVs for 24 h. Culture supernatants were collected and supplemented with 1mM N-ethylmaleimide to block the activity of deubiquitinases. The culture supernatant were used to purify sEVs by differential ultracentrifugation, and sEV were lysed on ice in IP buffer (50 mM Tris, pH8.0, 150 mM NaCl, 1% NP-40, supplied with protease inhibitors (1 mM AEBSF, 0.8 mM Aprotinin, 15 mM E-64, 10 mg/mL Bestatin, 10 mg/mL Pepstatin A and 10 mg/mL Leupeptin) and 1mM N-ethylmaleimide). SMO^FLAG^ was captured by incubation with M2 agarose beads (A2220-1ML, Sigma-Millipore) for 2 h at 4□. After three washes with IP buffer, bound proteins were eluted by incubation in LDS sample buffer for 10 min at 37 °C.

### Deubiquitinase treatment

*Ift27^-/-^* IMCD3 [SMO-FLAG] cells were cultured to near confluency and then serum-starved to induce ciliation and release of ciliary EVs for 24 h. The culture supernatant were used to purify sEVs by differential ultracentrifugation, sEV samples were lysed on ice in DUB buffer (50 mM Tris-HCl, pH 7.5, 1% NP-40, 25 mM KCl, 5 mM MgCl_2_, 1 mM DTT), and lysed sEVs were treated with 2 μM USP2 or 2 μM AMSH ^43,44^ for 1 h at 37 °C. Samples were then mixed with LDS sample buffer and incubated for 10 min at 37 °C.

### Immunoblotting

Cells were scraped from the plate into lysis buffer (0.5% v/v Triton X-100, 0.1% w/v SDS, 0.3 M NaCl, 1 mM EDTA, 1 mM EGTA, 25 mM Tris-HCl pH 7.5) supplemented with protease inhibitors (1 mM AEBSF, 0.8 mM Aprotinin, 15 mM E-64, 10 mg/mL Bestatin, 10 mg/mL Pepstatin A and 10 mg/mL Leupeptin), and lysates were added combined with LDS sample buffer (247mM Tris-HCl, 2% w/v LDS, 10% w/v Glycerol, 0.51 mM EDTA, 0.22 mM Coomassie G250, 0.175 mM Phenol Red, pH 8.5). EV samples were directly lysed with LDS sample buffer. The samples were supplemented with 20 mM DTT (except for samples to be probed for CD9 and GP96) and incubated at 37 °C for 10 min. Samples were loaded onto 4-12% SDS-PAGE gels (GenScript or Invitrogen), and transferred onto PVDF membranes using a Criterion Blotter wet transfer cell (Bio-Rad).

Membranes were blocked in 10 % SeaBlock (Thermo Scientific) in TBST or 5% milk (Bio-Rad) in TBST for 1 h at room temperature, then incubated with primary antibody overnight at 4 °C. After incubation with HRP-conjugated secondary antibody or HRP–Protein A (Jackson ImmunoResearch), blots were developed with ECL or Atto chemiluminescence detection kits (Pierce), and imaged on a Chemidoc Touch system (Bio-Rad).

### Anti-SMO antibody

A polyclonal rabbit antiserum against mouse SMO was generated by Prosci (Poway, CA). Rabbits were immunized with a GST fusion to a SMO protein fragment spanning aa. 560-793 that corresponds to the cytoplasmic C-tail of SMO exclusive of helix 8. Antibodies were affinity purified from the terminal antiserum on a column with a hexahistidine fusion to SMO[560-793] covalently immobilized to Sepharose beads (CNBr-sepharose, Cytiva).

### APEX labeling and proteomics of cilia-derived EVs

APEX labeling was performed as described previously ^25,26^. IMCD3 *Ift27^-/-^* cells expressing cilia-APEX2 (NPHP3[1-203]-APEX2) or control-APEX2 (NPHP3[1-203]G2A-APEX2) were seeded at high density and ciliation was induced by serum-starvation. The cells were pre-incubated with 0.5mM biotin-phenol for 120 min, labeled for 3 min by the addition of 1mM H_2_O_2_, and the reaction was determined by washing the cells in quenching buffer (1x PBS supplemented with 10 mM sodium ascorbate, 10 mM sodium azide and 5 mM Trolox). Cells were returned to DMEM/F12 supplemented with SAG and EV production was allowed to proceed by returning cells to 37 °C for 2 h. The culture supernatant was collected and subjected to sEV purification. The sEV fraction was lysed and the biotinylated proteins captured on streptavidin-sepharose beads. The captured proteins were digested with trypsin on beads and identified and quantitated by LC-MS^2^. Data was analyzed using Preview and Byonic (ProteinMetrics) for spectral counting-based quantitation. Missing spectral counts (SpC = 0) were imputed with 0.2 to calculate cilia-APEX2/control-APEX2 ratios. Proteins identified with SpCs < 4 (from both samples) were excluded from the list. Candidate proteins of the Cilia-APEX2 sEV proteome were defined as having cilia-APEX2/control-APEX2 ratios ≥ 4.

### TMS-MS of sEVs

#### SP3 bead binding and digestion

Purified extracellular vesicles were processed for proteomic analysis using a modified SP3 bead protocol^40^. Lysates were diluted to a final volume of 250 μL with 50 mM HEPES, 50 mM NaCl. Freshly made 1 M DDT in water was added to the samples to a final concentration of 5 mM, samples were placed on a shaker at 1,000 rpm for 30 min at 60 °C. After cooling to room temperature, the reduced proteins were alkylated with freshly prepared 400 mM iodoacetamide in 50 mM ammonium bicarbonate at a final concentration of 20 mM, the reaction proceeded in the dark for 30 min. Alkylation was quenched by the addition of 1 M DTT in water to a final concentration of 50 mM at room temperature for 15 minutes. A stock of 50 µg/µl SP3 beads in HPLC grade water containing hydrophobic and hydrophilic Sera-Mag SpeedBeads at a ratio of 1:1 (w/w) was prepared as described^40^. To each sample was added 10 μL of the 50 µg/µL SP3 magnetic beads, followed by the addition of 250 μL ethanol. Samples were gently vortexed to homogenize, and the mixture incubated at 24 °C for 5 minutes at 1,000 rpm. Using a magnetic rack, the supernatant was removed from the beads. The protein bound beads were washed, and the supernatant was removed a total of three times using 250 µl of 80% ethanol.

Proteins were then digested on bead, using 50 μL of digestion buffer (200 mM EPPS pH 8.5, 2% acetonitrile (v/v)), followed by LysC (2 mg/mL) at an enzyme-to-substrate ratio of 1:50 for 3 h at 37 °C. An additional 50 μL of digestion buffer was added along with trypsin (2 mg/mL) at an enzyme-to-substrate ratio of 1:100 and samples digested for 7 h at 37 °C. Beads were removed with a magnetic rack and clear supernatants transferred to new tubes.

### TMT labeling

TMT labeling was performed directly on digested samples. Acetonitrile 30% (v/v) was added to each sample followed by TMTpro 16plex reagent. The reaction was performed at room temperature with vortexing every 10 minutes for 1 h. To determine the TMT labeling efficiency and ratio check a pooled 12plex sample was analyzed by MS3 and confirmed labeling was >95%. The labeling reaction was quenched by the addition of hydroxylamine 50% aqueous solution to a final concentration of 0.5% (v/v), and incubated for 15 minutes. The quenched TMT samples were then acidified with formic acid to a pH below 3.0 and samples pooled together to form the 12plex. The solvent was evaporated under reduced pressure to near completion for subsequent bench-top fractionation by alkaline reversed phase chromatography (Pierce High pH Reversed Phase Peptide Fractionation Kit). Peptides were fractionated into 12 fractions by eluting with a 12-step gradient of acetonitrile, 10%, 12.5%, 15%, 17.5%, 20%, 25%, 30%, 35%, 40%, 50%, 65% and 80% acetonitrile. The acetonitrile fractions were combined using the following pattern 1^st^ + 7^th^, 2^nd^ + 8^th^, 3^rd^ + 9^th^, 4^th^ + 10^th^, 5^th^ + 11^th^, 6^th^ + 12^th^, which resulted in the final fractionated samples. The final samples were dried to completion followed by sample clean-up by Stage tip.

### Mass spectrometry

Peptides were separated using an EASY-nLC 1200 HPLC (ThermoFisher Scientific) on a capillary column (30-40 cm length, internal diameter of 100 μm) packed with 2.6 μm Accucore beads (Thermo Fisher Scientific) and heated to 60 °C, prior to analysis on an Orbitrap Fusion Lumos Tribrid MS. Peptides of each fraction separated using a flow rate of 450 nL/min with a gradient of 5-100% Buffer B over 240 min with Buffer A comprising 0.125% FA and Buffer B comprising 95% acetonitrile, 0.125% FA. All data were collected using a multi-notch MS3 TMT method ^45^. MS scans were performed in the Orbitrap over a scan range of 400-1400 m/z with dynamic exclusion of 2 minutes. Turbo rate (MS2) scans were performed in the Ion Trap with collision-induced dissociation energy (CID) of 35% and maximum injection times of up to 400 ms. TMT reporter ions were quantified using synchronous precursor selection (SPS-MS3) in the Orbitrap with a scan range of 100-1000 m/z after fragmentation by high-energy collision-induced dissociation (HCD) of 55%. Orbitrap resolution was set to 50,000 with varying injection times of up to a maximum of 600 ms. Further details on LC and MS parameters and settings used were described recently^46^.

### MS data analysis

Peptides were searched with a SEQUEST (v.28, rev. 12) based software against a size-sorted forward and reverse peptides sequences database of the mus musculus proteome (Uniprot 03/2021) with common contaminant proteins added. Searches used a mass tolerance of 20 ppm for precursors and a fragment ion tolerance of 0.9 Da. Maximally 2 missed cleavages per peptide were allowed for the searches. Oxidized methionine residues (+15.9949 Da) and were searched dynamically, while static modifications were used for cysteines alkylated with iodoacetamide (+57.0215 Da) along with TMT 16plex reagent (+229.1629 Da) on peptide N-termial and lysine.

A target decoy database strategy was applied and a false discovery rate (FDR) of 1% was set for peptide-spectrum matches (PSM) following filtering by linear discriminant analysis (LDA). A following FDR for final collapsed proteins was 1%. MS1 data were calibrated post search and searched again. Quantitative information on peptides was derived from MS3 spectra, TMT signal to noise quantification required an MS2 isolation specificity of ≥70%, and a summed signal to noise of ≥200 for all TMT channels for any given peptide. Details of the TMT intensity quantification method and further search parameters applied were described previously^46,47^.

### Statistical analysis

All graphs for quantification were generated, and statistical analyses were performed using Prism 9.0.0 (GraphPad). Data in **Figs. 3B, 4D and 5C** were analyzed using unpaired *t*-tests.

