## Supplement for "K63-linked ubiquitin chains mark inactive Smoothened for packaging into ciliary extracellular vesicles"

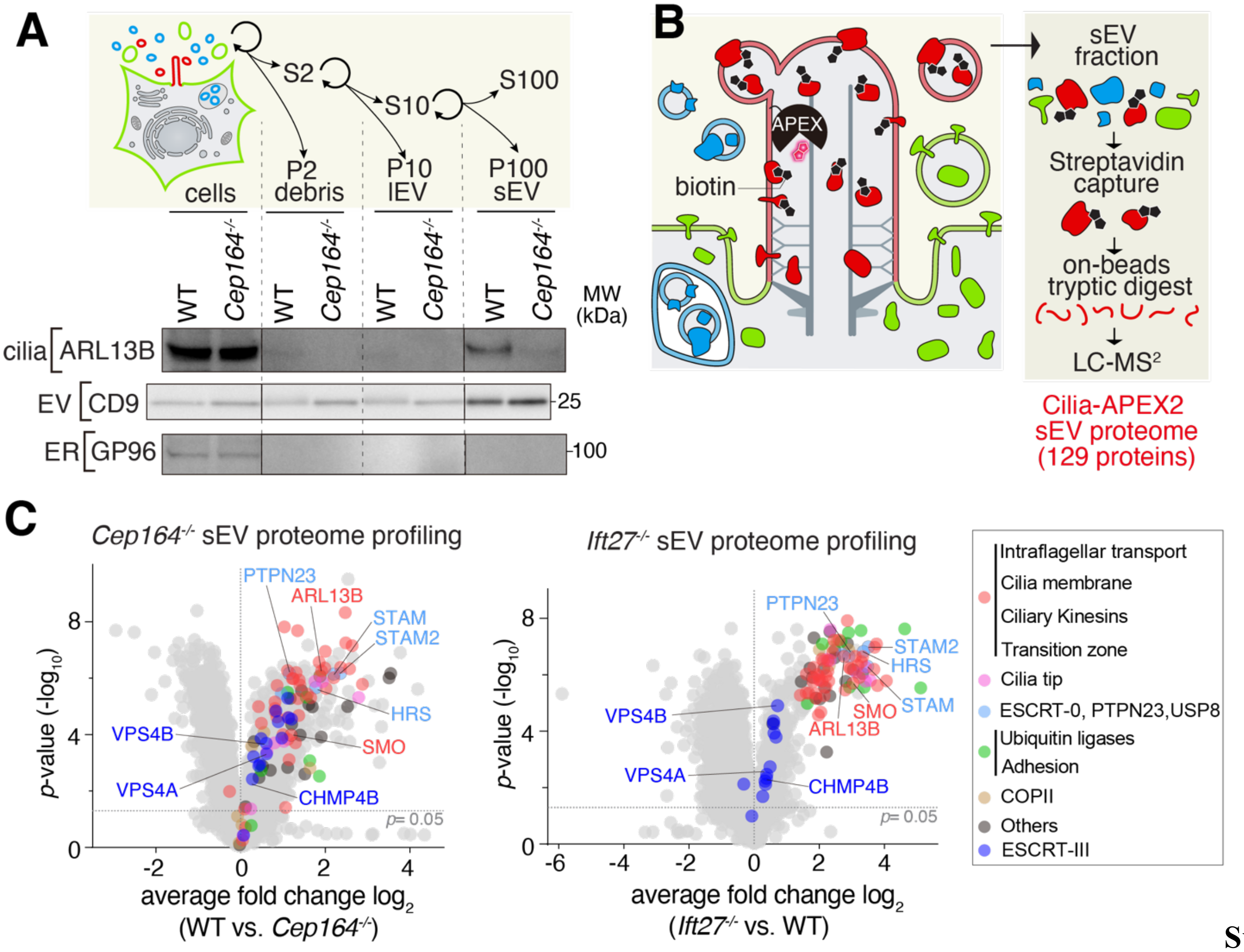


**Supplemental**

**Supplemental Figure 1. EV purification and proteomics of ciliary EVs.**

**A.** Fractionation of culture supernatants into EV fractions. IMCD3 cell supernatants were collected and EVs were purified by differential ultracentrifugation as described in the methods. The pellets of the 2,000 x *g* (P2), 10,000 x *g* (P10), and 100,000 x *g* (P100) centrifugation steps were collected and immunoblotted alongside the cell lysates for the cilium marker ARL13B, the generic EV marker CD9, and the ER marker GP96. 10 µg of cell lysates (corresponding to 0.27-0.36% of the total sample) were loaded for all immunoblots. For ARL13B detection, 15% of the P2 and P10 fractions and 50% of the P100 fractions were loaded. For CD9 and GP96 detection, 15% of the P2 and P10 fractions and 10% of the P100 fractions were loaded.

**B**. Strategy for proteomics of ciliary sEVs via Cilia-APEX2. Ciliary proteins were biotinylated by treating IMCD3 *Ift27*$^{-/-}$ cells expressing cilia-APEX2 (or-control APEX2) with biotin-phenol and $H_2O_2$. The labeling reaction was terminated by washing cells in quenching buffer and EV production was allowed to proceed for 2 h at 37 ºC in the presence of SAG. sEVs purified from the culture supernatant were lysed, biotinylated proteins captured on streptavidin beads, and captured proteins digested with trypsin on beads, before peptide identification by LC-MS$^2$.

**C**. Volcano plots comparing the proteomes of WT vs. *Cep164*$^{-/-}$ sEVs or *Ift27*$^{-/-}$ vs. WT sEVs. Statistical *p* values for 2,640 quantified proteins were plotted against the TMT ratios of WT samples versus *Cep164*$^{-/-}$ samples (left) or *Ift27*$^{-/-}$ samples versus WT samples (right). Proteins of the ciliary sEV cluster and ESCRT-III are highlighted by colored dots according to their subgroup categories as indicated, and other proteins are shown in gray. Nearly all proteins from the ciliary sEV cluster were significantly enriched in WT vs. *Cep164*$^{-/-}$ sEV, and all proteins from the ciliary sEV cluster were significantly enriched in *Ift27*$^{-/-}$ vs. WT sEV.

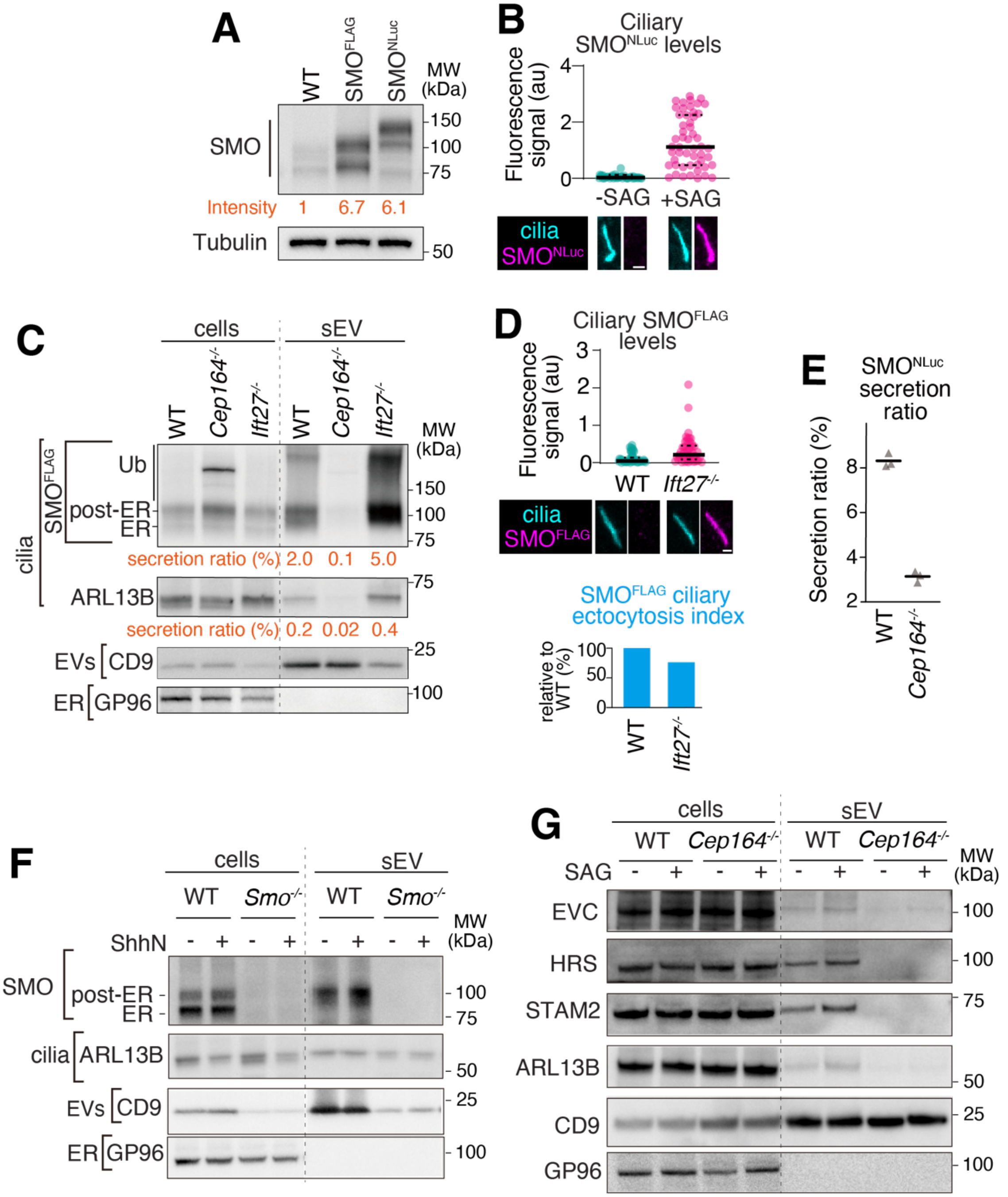


**Supplemental Figure 2. Regulation of SMO secretion by signaling.**

**A**. Comparison of the expression levels of SMO$^{FLAG}$ and SMO$^{NLuc}$ to endogenous SMO. Equal amounts of cell lysates from IMCD3 cells, IMCD3-[SMO-FLAG], and IMCD3-[SMO-FLAG-NLuc] cells were immunoblotted for SMO and α-tubulin. The relative intensities of the SMO bands normalized to α-tubulin were quantified and are shown below the immunoblots.

**B.** Ciliary enrichment of SMO$^{NLuc}$ in response to SAG. Near-confluent and serum-starved IMCD3-[SMO-FLAG-NLuc] cells were treated with or without 200nM SAG for 20 h before staining for FLAG and acetylated tubulin (cilia). The ciliary fluorescence intensity in the FLAG channel was plotted in a dot plot. In this and all dot plots, the thick bar indicates the median and the dotted lines the first and third quartiles. $N$ = 50 cilia analyzed per condition; au, arbitrary units. Representative images under the graph. Scale bar, 1 µm.

**C-D.** Secretion of SMO$^{FLAG}$ in WT, cilia-less and retrieval-defective cells. Near-confluent WT, *Cep164*$^{-/-}$, and *Ift27*$^{-/-}$ IMCD3-[SMO-FLAG] cells stably were serum starved and EV secretion allowed to proceed for 20 h. **C.** sEVs were prepared from the culture supernatants, and cell lysates and sEV fractions were immunoblotted for FLAG, ARL13B, CD9, and GP96. 10 µg of cell lysates (corresponding to 0.18-0.2% of the total sample) were

loaded for all immunoblots. For SMO and ARL13B detection, 30% of the P100 fractions were loaded. For CD9 and GP96 detection, 10% of the P100 fractions were loaded. The secretion ratios of $SMO^{FLAG}$ and ARL13B were quantified and are shown below the immunoblots. **D.** Cells were stained for FLAG and acetylated tubulin (cilia) and the ciliary fluorescence intensity in the FLAG channel was plotted in a dot plot. $N = 50$ cilia analyzed per condition. Representative cilia images shown below the graph. Scale bar, 1 µm. The ciliary ectocytosis indices of $SMO^{FLAG}$ are shown in a bar graph.
**E.** Requirement for cilia in $SMO^{NLuc}$ secretion. Near-confluent WT and *Cep164*$^{-/-}$ IMCD3-[SMO-FLAG-NLuc] cells were serum-starved for 20 h. The levels of $SMO^{NLuc}$ in culture supernatants and in cells were measured using the NanoLuc assay. Each grey triangle represents the $SMO^{NLuc}$ secretion ratio from one experiment. $N = 3$ independent experiments.
**F.** Validation of the anti-SMO antibody. WT and *Smo*$^{-/-}$ NIH-3T3 cells and sEV fractions treated as in **Fig. 2E** were immunoprobed with the anti-SMO antibody.
**G.** Regulation of EVC, HRS, STAM2, ARL13B secretion by SAG. Near-confluent IMCD3 WT and *Cep164*$^{-/-}$ cells were serum-starved and treated with or without 200 nM SAG for 16 h. sEVs were prepared from the culture supernatants, and cell lysates and sEV fractions were immunoblotted for EVC, HRS, STAM2, ARL13B, CD9, and GP96. 0.4% of the total cell lysates were loaded for all immunoblots. For ARL13B and EVC detection, 20% of the P100 fractions were loaded. For HRS and STAM2 detection, 40% of the P100 fractions were loaded. For CD9 and GP96 detection, 10% of the P100 fractions were loaded.

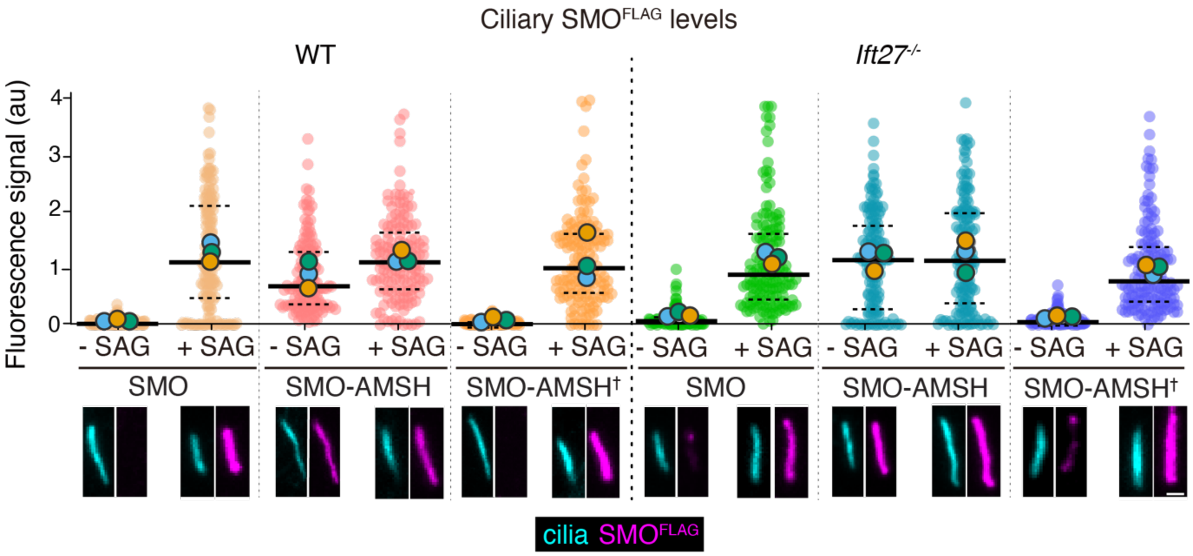


**Supplemental Figure 3. Effect of forced deubiquitination of the ciliary levels of SMO.**

This panel adds the SAG-treated condition to **Fig. 5C**, where only ciliary SMO$^{FLAG}$ of untreated cells is shown. WT and *Ift27*$^{-/-}$ IMCD3 cells stably expressing the indicated SMO fusions were treated with or without 200 nM SAG for 20 h. Cells were stained for FLAG and acetylated tubulin (cilia) and the ciliary fluorescence intensity in the FLAG channel was plotted in a dot plot. *N* = 150-151 cilia from 3 independent experiments analyzed per condition; au, arbitrary units. Representative cilia shown below the graph. Scale bar, 1 µm.

**Table S1**

| **Chemicals, Peptides, and Recombinant Proteins** | | |
|---|---|---|
| Rabbit polyclonal anti-EVC | Sigma-Aldrich | HPA008703 |
| Cyclopamine | Selleckchem | S1146 |
| SAG (Smoothened Agonist) | Selleckchem | S7092 |
| SANT-1 | American Peptide Company | AP68-1-10A |
| Ponasterone A | Sigma-Aldrich | P3490 |
| Dual-Luciferase® Reporter Assay System | Promega | E1960 |
| HEK 293T EcR-Shh | ATCC | CRL-2782 |

**Data S1.**
Proteomics of Cilia-APEX2 sEVs.

**Data S2**
Proteomic profiling of sEVs from WT, *Ift27*$^{-/-}$ and *Cep164*$^{-/-}$ cells.
